# FAM162A is Crucial for Mitochondrial Structure, Dynamics, and Bioenergetics, Driving Cellular Protection and Longevity

**DOI:** 10.1101/2024.12.26.630390

**Authors:** Andrea Matamoros, Juan Pablo Soffia, Marcelo Muñoz, Michael Maturana, Alvaro Gonzalez-Ibañez, Gabriela Gomez-Lillo, Cesar Astorga, Lina M. Ruiz, Ramón Jorquera, Alvaro A. Elorza

## Abstract

**Introduction:** FAM162A is a mitochondrial protein evolutionarily conserved across taxa and ubiquitously expressed in various tissues. It is known for its role in hypoxia-induced apoptosis. However, paradoxically, FAM162A is overexpressed in cancer, where its pro-apoptotic function seems overridden, suggesting an alternative role associated with mitochondrial function and cell survival. Additionally, its precise localization and topology remain controversial.

**Objectives:** To assess the role of FAM162A in mitochondrial structure, dynamics, and bioenergetics and its impact on cell viability, while establishing its precise localization, orientation, and topology. Additionally, to generate a transgenic Drosophila model overexpressing human FAM162A to evaluate its effects on organismal survival under normal and stress conditions.

**Methods:** Localization, orientation, and topology were determined by protease protection assays in COS7 cells. Loss-and gain-of-function experiments were performed to assess mitochondrial function and turnover by confocal microscopy, immunoblots and Seahorse technology. A transgenic Drosophila model overexpressing human FAM162A was generated to evaluate organismal survival under normal and stress conditions.

**Results:** FAM162A is essential for maintaining mitochondrial ultrastructure and bioenergetics, thereby influencing cell viability and stress resistance. Localization studies revealed that FAM162A resides predominantly in the inner mitochondrial membrane, particularly within the cristae, where it modulates the fusion protein OPA1. Transgenic Drosophila overexpressing human FAM162A exhibited increased lifespan and locomotor activity under both normal and heat stress conditions.

**Conclusion:** FAM162A emerges as a crucial player in maintaining mitochondrial integrity and bioenergetics. Its functional role, potentially mediated through interaction with OPA1, impacts mitochondrial health, stress resistance, cellular viability, and organismal longevity.

## Introduction

Mitochondria are essential cellular organelles, often referred to as the powerhouses of the cell, responsible for meeting the cell’s ATP demands through oxidative phosphorylation. Beyond their well-established role in energy production, mitochondria are now recognized as pivotal metabolic hubs. They integrate various intracellular signaling pathways to coordinate and execute cellular responses, thereby enabling the adaptation of cell metabolism to both external and internal environmental changes. This capability positions mitochondria at the forefront of cellular fate determination, stress response signaling, and metabolic reprogramming, influencing diverse cellular processes such as apoptosis, autophagy, cell proliferation and differentiation, aging, and age-related diseases.

Mitochondria distribute as a complex network that remodels thought fusion and fission events to respond to metabolic demands as well as to adapt to stressful cellular environments. These processes known as mitochondrial dynamics are not only responsible of mitochondrial size and morphology, but also and most importantly, of bioenergetics outputs. These include the respiratory rate, energy expenditure and ATP synthesis as well as apoptosis and the segregation and elimination of dysfunctional mitochondrial units by mitophagy (Twig et al., 2008; Wikstrom et al., 2014; Westermann, 2010; Sheridan and Martin, 2010). Mitochondrial fission relies on the cytosolic protein DRP1 which translocates to mitochondria and binds the outer mitochondrial membrane (OMM)- receptor proteins FIS1, MFF, MiD49 and MiD51 (Losón et al., 2013; Gandre-Babbe and Bliek, 2008; Youle and Bliek, 2012; Held and Houtkooper, 2015). DRP1 mediates fission by forming a multimeric ring around the OMM. Mitochondrial fission plays a crucial role in facilitating mitochondrial quality control (Pickles et al., 2018), mtDNA replication and distribution (Ota et al., 2020), organelle division during the cell cycle (Otera et al., 2013) and apoptosis among other cellular events. Mutations in DRP1 and MFF are associated with developmental disorders due to defective OXPHOS and overall mitochondrial dysfunction (Robertson et al., 2023). Mitochondrial fusion is a critical process for maintaining the integrity and functionality of mitochondria. It involves the mixing and redistribution of mitochondrial contents, ensuring a uniform population of mitochondria. Fusion also allows for genetic material exchange, which helps repair damaged mitochondrial DNA. Key players in mitochondrial fusion are the Mitofusins (MFN1 and MFN2) and OPA1. MFN1 and MFN2 facilitate the OMM fusion, while OPA1 regulates the inner mitochondrial membrane (IMM) fusion, influencing cristae remodeling and shape (Chen et al., 2003; Song et al., 2009; Hu et al., 2020). Disruption of fusion can lead to the accumulation of dysfunctional mitochondria and increased ROS production. Mutations in MFN2 and OPA1 are linked to Charcot-Marie-Tooth disease type 2A and dominant optic atrophy, respectively (Züchner et al., 2004; Zanna et al., 2008). Loss of fusion results in decreased mitochondrial DNA content, membrane potential loss, and impaired respiratory chain function (Jang and Javadov, 2020; Xin et al., 2021).

Maintaining mitochondrial health is crucial for cellular function. Conversely, dysfunctional mitochondria, characterized by unbalanced MtDy, impaired bioenergetics and disrupted ultrastructure, can lead to a cascade of detrimental effects, including increased oxidative stress, defective ATP production, and altered metabolic signaling. These disruptions are implicated in a range of pathological conditions, from metabolic disorders to neurodegenerative diseases and cancer.

FAM162A, also known as HGTD-P, is a mitochondrial protein discovered in 2004 (Lee et al., 2004) as a target of the transcription factor HIF-1α and it is highly conserved among evolutionary distant taxa and ubiquitously expressed in many tissues. At the organism level, FAM162A mRNA displays higher expression in colon, esophagus, heart, kidney, and liver (Fagerberg et al., 2014). FAM162A has been described to participate in hypoxia-induced apoptosis through the binding to the mitochondrial voltage-dependent anion channel (VDAC) to stimulate the opening of the mitochondrial permeability transition pore (mPTP). However, FAM162A has also been reported to be overexpressed in several types of cancer, but far from inducing apoptosis and cell death, cancer cells increased their proliferation and migration capacities (Tang et al., 2009). These contradictory antecedents suggest an alternative function from apoptosis for FAM162A which has not been described so far. Furthermore, FAM162A was initially proposed to contain two transmembrane segments based on protein deletion studies and mitochondrial import observations (Lee et al., 2004). However, a bioinformatic prediction using TOPCONS (https://topcons.cbr.su.se/), published by Lee, S.-H. et al (2020)(Lee et al., 2020), suggested a single transmembrane segment located in the OMM between amino acids 102 and 122. According to this prediction, the N-terminus would face the inter membrane space while the C-terminus would face the cytosol (Lee et al., 2020). Thus, the exact localization of FAM162A within mitochondria, its orientation, and topology has not been afforded empirically and, therefore, remains unconfirmed.

This study elucidates the role of FAM162A in mitochondrial function, highlighting its subcellular localization and impact on cellular function and the entire organism. By employing loss-and gain-of-function experiments in COS7 cells, along with transgenic Drosophila as an organismal model, we demonstrate that FAM162A localizes predominantly to the inner mitochondrial membrane, particularly within the cristae. Our findings reveal that FAM162A plays a crucial role in mitochondrial bioenergetics and dynamics, contributing to mitochondrial turnover and offering protection against cellular stress. Notably, overexpression of human FAM162A in transgenic Drosophila extended life expectancy by 25% under normal conditions and 40% under heat stress. These results uncover significant new roles for FAM162A, advancing our understanding of its function in mitochondrial regulation and organismal resilience.

## Materials and Methods

### Cell culture and cell transfection

COS7 cells were cultured in Dulbecco’s Modified Eagle’s Medium (DMEM) with 1.0 g/L D- Glucose, 110mg/L Sodium Pyruvate (01-050-1A, Biological Industries), supplemented with 2mM L- Glutamine (25-005-C1, Corning), 1% penicillin/streptomycin (03-031-1, Biological Industries), and 10% fetal bovine serum (04-127-1A, Biological Industries). Cells were incubated at 37°C with 5% CO2. Testing for mycoplasma contamination was performed once a month, being negative throughout this study. Cell transfection was performed with Lipofectamine 2000 (Invitrogen) following the manufacturer’s instructions.

### Molecular tools and reagents

To assess the role of FAM162A, three siRNA for FAM162A were designed (Table 1) and cloned into the vector pLVCTH (Tronolab) to generate the pLVCTH-siFAM162A-GFP constructs for gene silencing. In addition, the construct pcDNA3.1_FAM162A_Myc/His/A for FAM162A overexpression (FAM162A-OE), was sent to synthesize to GenScript (New Jersey, USA). The following reagents were utilized throughout this research: FCCP (Carbonyl cyanide-*p*-trifluoromethoxyphenylhydrazone ab120081), Rotenone (R8875, Sigma), Paraquat (Methyl viologen 856177, Sigma), Oligomycin (O4876, Sigma), Antimycin A (A8674, Sigma).

**Table 1:**
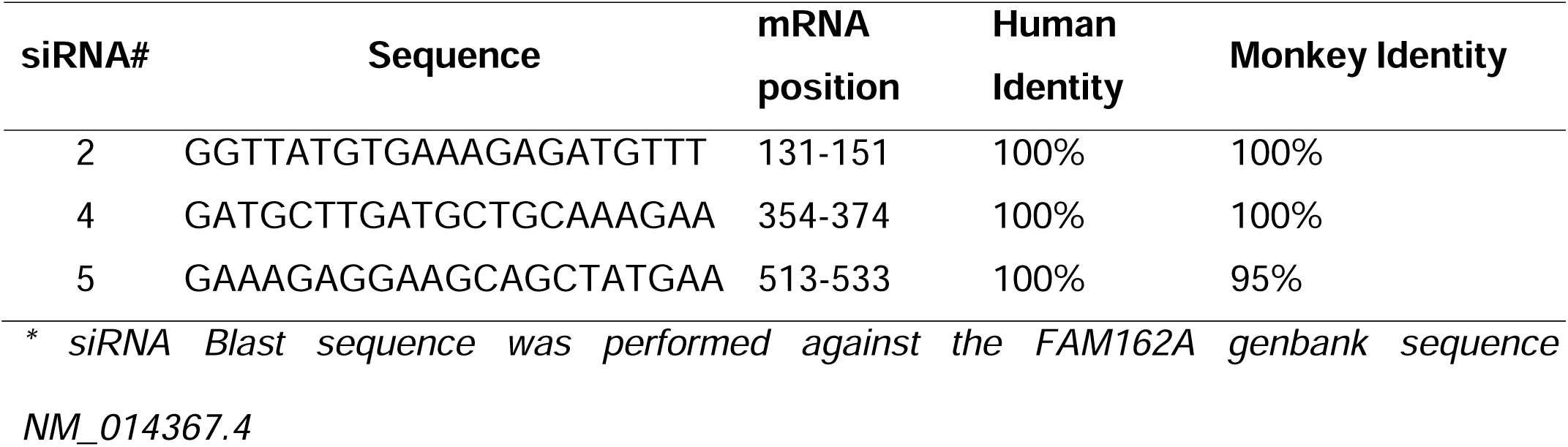
Three siRNA targeting human *Fam162A* transcripts were designed and aligned against human and green monkey *fam162a* gene.

### Proteinase K protection Assays

The protease protection assay was selected to establish the location and orientation of FAM162A which was evaluated by both live cell confocal microscopy and Western Blot (Lorenz et al., 2006). For confocal microscopy, the constructs FAM162A_pcDNA3.1(+)-N-eGFP (FAM-N-GFP) and FAM162A_pcDNA3.1(+)-C-eGFP (FAM-C-GFP) which contain GFP fused either to the N-or C- - terminus of FAM162A, were used. Additionally, the OMP25_mCherry plasmid was used as a control marker for outer mitochondrial membrane, having the mCherry protein facing the intermembrane space (Nemoto and Camilli, 1999); the pmMitoTurquoise plasmid, as a control marker for the inner mitochondrial membrane, having the turquoise protein facing the mitochondrial matrix (Zong et al., 2018); and the pEGFP-N1 (Clontech) as a control marker for the cytosolic compartment. For live confocal microscopy experiments, cells were transfected and 24 hrs. later, placed in the Chamlide chamber (Live cell instrument, South Korea) to be sequentially exposed to 25µg/ml or 400µg/ml Digitonin for 3 min and then followed by 50µg/ml Proteinase K. Confocal images were acquired pre-digitonin (time 0), and post digitonin (time 180). Then, Proteinase K was added, and images were taken at 210, 240 and 300s. Images were analyzed with the Fiji software. For the Western blot experiments, it was used the construct FAM162A_pcDNA3.1(+)-myc-His A which contain c-myc tag fused to the C-terminus of FAM162A. Additionally, GAPDH was used as a free cytosolic protein; TOM20, that is inserted in the OMM but having most of its protein body in the citosol, was used as a membrane-bound cytosolic protein control; and the SDH2A subunit, as a mitochondrial matrix control. Cells were transfected with FAM162A-c-myc and 24 hrs. post-transfection, treated with Digitonin and Proteinase K as described above.

### Western Blot

Cell pellets were homogenized in NP40 lysis buffer (150mM NaCl, 1% NP-40, 50mM Tris pH 8.0, 0.1% SDS plus the protease inhibitors cocktail (Cat. # P8340, Sigma-Aldrich) and PMSF (Cat #36978, Thermo Fischer Scientific). Total proteins (16µg) were fractionated in a 15% 2,2,2- trichloroethanol-containing polyacrylamide gels and transferred to 0,2 µm PVDF membranes. The incorporation of trichloroethanol enables fluorescent visualization of proteins under UV light exposure serving as a protein loading control. Blots were blocked in 5% non-fat dry milk in TBS-T buffer (Tris-buffered saline, pH 7.4, 0.2% Tween 20) during 1hr, and then incubated with the primary antibodies (Table 2) in TBS-T overnight at 4 °C. After three washes with TBS-T for 5 min each, blots were incubated with the HRP-conjugated secondary antibody (Table 2) in blocking solution for 1 hour. Finally, membranes were treated with the Super Signal West Femto (Cat # 34094, Thermo Fischer Scientific) and revealed with the Alliance Q9 Advanced UVITEC instrument. Band densitometric analysis was performed with the NINE ALLIANCE UVITEC software.

**Table 2:** Antibodies used in this study and their dilutions for Western Blot (WB)

| Antigen | Code | Source | Type | WB |
| --- | --- | --- | --- | --- |
| MFN1 | ab57602 | abcam | mouse | 1:1000 |
| MFN2 | ab50843 | abcam | rabbit | 1:1000 |
| OPA1 | ab42364 | abcam | rabbit | 1:1000 |
| OXPHOS | ab110413 | abcam | mouse | 1:1000 |
| c-Myc-tag | 2278 | Cell Signaling | rabbit | 1:1000 |
| DRP1 Ser616 | 3455S | Cell Signaling | rabbit | 1:1000 |
| GAPDH | 2118 | Cell signaling | rabbit | 1:2500 |
| Mouse IgG (H+L) HrP | 31432 | Invitrogen | goat | 1:10000 |
| Rabbit IgG (H+L) HrP | 31460 | Invitrogen | goat | 1:10000 |
| DRP1 | sc271583 | Santa Cruz | rabbit | 1:500 |
| HGTD-P (FAM162A) | sc-514243 | Santa Cruz | rabbit | 1:300 |
| SDHA | sc-390381 | Santa Cruz | mouse | 1:500 |
| TOM-20 | sc-17764 | Santa Cruz | mouse | 1:500 |
| FIS1 | TF22142 | Thermo Fisher | rabbit | 1:1000 |

### Mitotimer Experiments

COS7 cells were seeded over 12 mm cover glasses and co-transfected with the pMitoTimer (Addgene #52659 gently donated by Dr. Roberta Gottlieb) and the pcDNA3.1_Myc/His/A backbone (EMPTY) or FAM162A-OE constructs. Then, 24 hours post-transfection, cells were treated with 100uM paraquat, a pro-oxidant. After the drug treatment, cells were fixed with 4% paraformaldehyde/sucrose for 3 min. After three washing steps, the coverslips were mounted on slides with 5uL Fluoromont-G (Invitrogen) and visualized in the confocal microscope LEICA TCS SP8. Fiji software was used to analyze and quantified the images.

### Transmission Electron Microscopy

FAM162 knockdown and control COS7 cells were fixed in 2.5% glutaraldehyde and 0.1 M cacodylate buffer, pH 7.2, for 6 hrs. at RT, and then rinsed with a 0.1 M cacodylate buffer, pH 7.2, for 18 h at 4 C. Right after, cells were immersed in 1% OsO_4_ for 90 min, rinsed with distilled water, and labeled with 1% uranyl acetate for 1 h. Cells were dehydrated in graded series of acetone to 100% and embedded in Epon resin. Polymerization reactions were done at 60 C for 24 h. Ultrathin sections (60– 70 *nm*) were obtained with the ultramicrotome Sorval MT-500, disposed in copper grids, and contrasted with 4% uranyl acetate in methanol for 2 min and lead citrate for 5 min. Grids were examined with a Philips Tecnai 12 electron microscope at 80 kV. Fiji software was used to analyze the images.

### Mitochondrial Bioenergetics

Cells were seeded in 3 cm glass bottom confocal plates for live cell confocal microscopy. Mitochondrial morphology and mitochondrial membrane potential were assessed by staining cells with 10nM TMRE (11560796, Invitrogen), in a non-quenching mode, for 30 min at 37°C. Then, live cell imaging was done under a confocal Olympus Fluoview 1000 microscope coupled to a temperature and CO2-controlled chamber (Chamlide TM IC, Live Cell Instrument, Inc). Single confocal images were acquired at 60x magnification. Images were analyzed with the Fiji software and the mean fluorescence intensity of whole cells was obtained for mitochondrial membrane potential. Mitochondrial morphology was assessed as described by Leonard (2015) (Leonard et al., 2015).

The oxygen consumption and the extracellular acidification rates was measured with the Seahorse XF24 or XF Pro Extracellular flux analyzer (Agilent) as described in Escalona et al. (2020) (Escalona et al., 2020). Cells were plated at 60.000 (XF24) or 20.000 (XF Pro) cells/well in the XF V7- PS (XF24) or XF Pro M plate (XF Pro) plates and then basal respiration, ATP-linked respiration, leaking, maximal respiration, and non-mitochondrial respiration were measured by the sequential addition of 5mM Glucose, 1μM oligomycin, 0.75μM FCCP and 1µM Rotenone/1µM Antimycin A. The ATP synthesis rate was measured with the Seahorse XF Real-Time ATP Rate Assay Kit (103592-100).

### Viability and cytotoxicity assays

The viability test was carried out using the MTT Cell Viability Assay (V13154 Invitrogen); and the cytotoxicity assay, using the LDH Cytotoxicity Assay LDH kit (88953 Thermoscientific), both according to manufacturer instructions.

### Transgenic Drosophila

Human FAM162A cDNA (Gene ID: 26355), optimized for Drosophila melanogaster codons, was cloned into the pUASTattB-5xUAS/Mini_Hsp70 vector (VectorBuilder Inc.). This UAS- hFAM162A construct was inserted into the Drosophila genome (BestGene, Inc.) to create UAS_FAM162A transgenic flies, enabling targeted expression. The UAS/GAL4 system was employed for ubiquitous FAM162A overexpression throughout the organism. Specifically, UAS_FAM162A flies (Control_UAS_FAM162A) were crossed with Tubulin-GAL4 flies (Control_Gal4) to generate UAS/GAL4_FAM162A_OE flies (hFAM162A_OE). The transgenic flies were maintained under standard laboratory conditions.

For lifespan analysis, 20 flies of each genotype (Control Gal4, Control UAS_FAM162A, and hFAM162A_OE) and sex (male and female) were housed in culture tubes with ad-libitum food. The experiment was conducted at 29°C, with fly survival monitored daily until natural death (approximately 40 days). Additionally, groups of 5 flies were subjected to heat stress at 40°C until death to assess stress resistance. Survival and locomotor activity were recorded on video to generate lifespan curves and assess velocity over time. The velocity parameter was adjusted by fly weight. Kaplan-Meier analysis was used to generate survival curves. All calculations, data analysis, statistics, and plotting were conducted in R Studio.

### Statistics

All calculations, data analyses, statistical tests, and plots were conducted using GraphPad Prism® version 8.0, Jamovi version 2.3.19, or R Studio version 4.4.1, with significance set at p ≤ 0.05. For comparisons of two mean values, the Student’s t-test was used for unpaired data. In Western blot experiments, each cell culture plate was considered as an experimental unit, with a minimum of three independent biological replicates per experiment. For microscopy experiments, individual GFP-positive cells were considered as experimental units. In Drosophila experiments, each fly was treated as an experimental unit. For fly survival analysis, Kaplan-Meier survival curves were generated.

## Results

### FAM162A is a highly conserved protein localized in the inner mitochondrial membrane

FAM162A is a hypoxia-induced proapoptotic protein, overexpressed in different types of cancer and is considered one of the 15-hypoxia markers. Blast analysis (Johnson et al., 2008) displayed that FAM162A is widely expressed between different taxa having a gene and protein homology ranging from 99% in monkeys to 50% in fish when compared to humans (Fig. 1A). FAM162A is then an evolutionary conserved gene which suggests an important role for cell biology. 3D protein structure modeled through AlphaFold 2.0 software (Jumper et al., 2021) displayed two transmembrane segments, an extended loop with a short alpha-helix domain, and a C-terminus alpha-helix structure (Fig. 1B).

**Figure 1:**
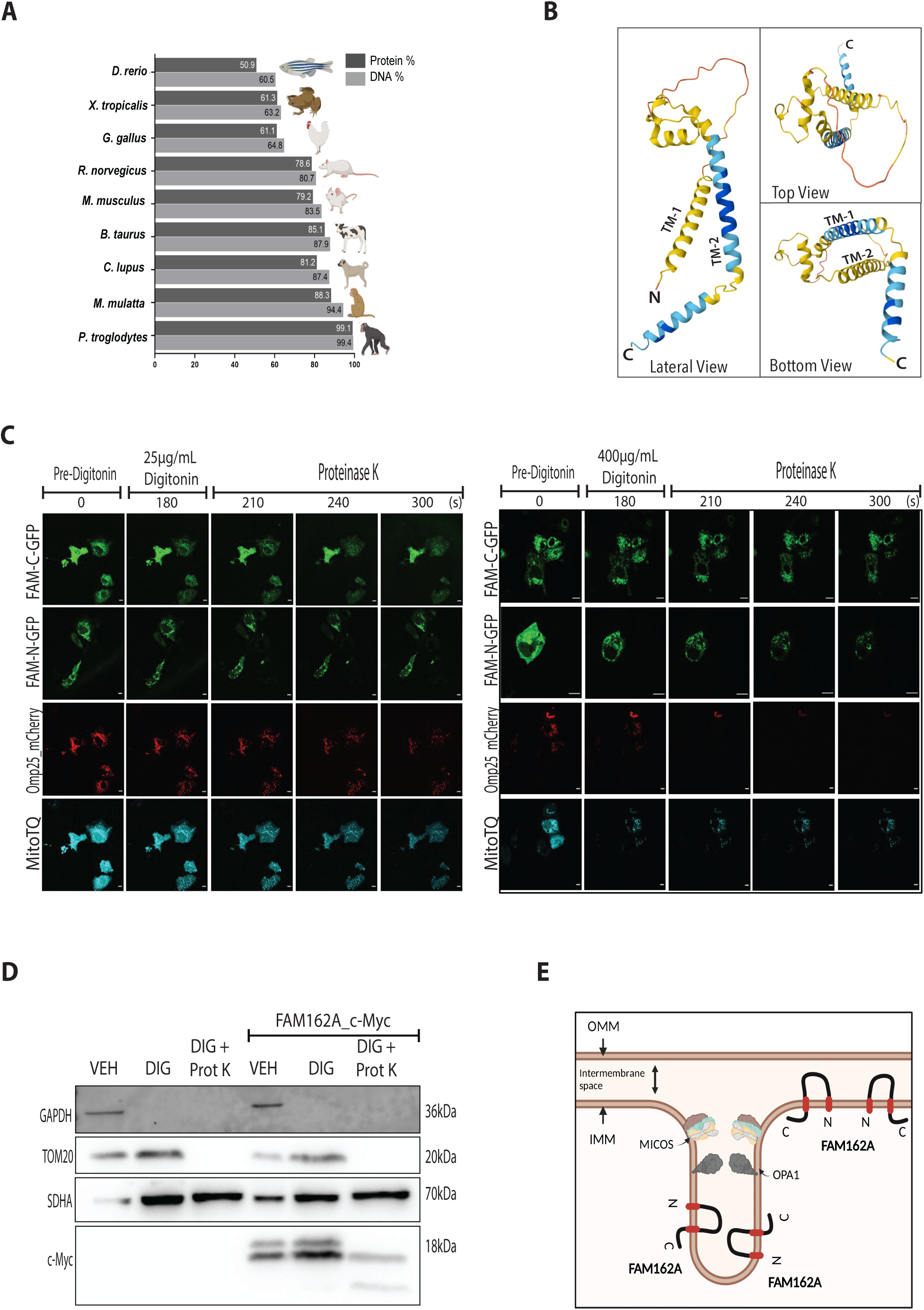
Comparative Conservation of FAM162A across Taxa, Predictive Structure, and Mitochondrial Localization (A) Pairwise Alignment. Scores from NCBI showing DNA and protein homology percentages of FAM162A in *Homo sapiens* versus other species, revealing that the FAM162A gene and protein are conserved among different taxa (HomoloGene, n.d). **(B) 3D Model of FAM162A Predicted by Alphafold 2.0.** Lateral, Top and Bottom views are shown. **The model reveals two putative transmembrane domains connected by an extended loop and a short alpha-helix domain at the C-terminus.** TM-1: Transmembrane Segment 1. TM-2: Transmembrane Segment 2 **(C) Fluorescence Protease Protection (FPP) Assay.** COS7 cells overexpressing FAM162A fused to GFP at either the N-terminus (FAM-N-GFP) or C-terminus (FAM-C-GFP) were subjected to FPP assay with 25 µg/ml (left panel) and 400 µg/ml (right panel) Digitonin. Controls included COS7 cells co-transfected with OMP25-mCherry (intermembrane space marker) and pmMitoTurquoise (mitochondrial matrix marker). Fluorescence was observed using live confocal microscopy. Under the 400 µg/ml Digitonin condition, OMP25-mCherry was degraded, while FAM-N-GFP and FAM-C-GFP fluorescence persisted similarly to MitoTQ, indicating localization within the inner mitochondrial membrane. Scale bars = 5 µm (*n*=3). **(D) Protease Protection Assay by Western Blot.** COS7 cells transfected with the FAM162A- OE_c-myc plasmid or empty vector were treated with 400 µg/ml Digitonin and Proteinase K. Immunoblotting was performed for c-myc (to detect FAM162A overexpression), GAPDH (free cytosolic protein), TOM20 (outer mitochondrial membrane protein exposed mostly to the cytosol), and SDHA (inner mitochondrial membrane protein facing the mitochondrial matrix) as controls. Two bands were observed for FAM162A-OE_c-myc. One, persisted similarly to SDHA, suggesting localization within the mitochondrial cristae and, therefore, fully protected from Proteinase K. The second, was partially degraded by Proteinase K suggesting localization in the inner boundary membrane (*n*=3). **(E) Cartoon model of FAM161A localization, orientation, and topology.** FAM162A predominantly localizes within the mitochondrial cristae, sheltered by the MICOS complex and OPA1 protein at cristae junction. A fraction of FAM162A is also found in the inner boundary membrane. FAM162A contains two transmembrane segments, with both N- and C- termini facing the mitochondrial matrix.

The exact localization, orientation, and topology of FAM162A within mitochondria has remained unconfirmed until now. Consequently, we decided to investigate it to gain more insights into its function using the protease protection assay. Briefly, we generated two constructs to overexpress FAM162A fused to GFP located either at the N-terminus (FAM-N-GFP) or C-terminus (FAM-C-GFP) in COS7 cells to perform a fluorescence protease protection assay (Fig.1C). As controls, we utilized COS7 cells co-transfected with the Omp25_mCherry and pmMitoTurquoise (MitoTQ) plasmids to label both intermembrane space and the mitochondrial matrix respectively. As shown in Figure 1C (*left panel*), when cells were treated with 25µg/ml Digitonin (to permeabilize only the plasma membrane) and Proteinase K, all fluorescent proteins remained intact, indicating protection from digestion. However, when cells were treated with 400µg/ml Digitonin (to permeabilize the plasma membrane and the OMM) and Proteinase K (Fig. 1C, *right panel*) the Omp25_mCherry fluorescence got extinct at 240-300s. Conversely, both FAM-N-GFP and FAM-C-GFP were protected similarly to MitoTQ, suggesting that both the N- and C-termini of FAM162A are located inside the mitochondrial matrix (Fig.1C, *right panel*).

To corroborate our findings, we also performed a standard protease protection assay analyzed by Western blot. COS7 cells were transfected with the FAM162A-OE_c-myc construct where the c-myc tag was placed at the C-terminus of FAM162A. Cells were treated with the Digitonin buffer alone (VEH), with 400 µg/ml Digitonin alone, and with Digitonin plus Proteinase K. Total proteins were isolated and immunoblotting was performed against c-myc (Fig. 1D). As localization controls, GAPDH expression level was used as a free-cytosolic protein; TOM20, as an OMM protein (with most amino acids facing the cytosol); and SDHA, as an inner mitochondrial membrane (IMM) protein (with most amino acids facing the mitochondrial matrix). The Western blot assay displayed two bands for FAM162A_c-myc under the vehicle and digitonin conditions, suggesting potential post-translational modifications. Interestingly, in the presence of proteinase K, the upper band was partially digested, generating a ∼10kDa band. Conversely, the lower band remained intact, indicating protection from digestion (Fig. 1D). Our results revealed intriguing findings, suggesting that FAM162A primarily localizes to the IMM, predominantly within the cristae membrane (CM), where it might be protected from digestion due to the sheltering effect of the cristae junction by MICOS complex and OPA1 protein (Fig. 1E). Furthermore, our data supported the presence of two transmembrane segments with both the N- and C-termini facing the mitochondrial matrix, and the loop residing within the cristae lumen (Fig. 1E). This localization and orientation of FAM162A in the IMM and CM provide insights into potential roles in maintaining cristae organization and the bioenergetic capacity of mitochondria.

### FAM162A is important for mitochondrial bioenergetics and cell viability

To assess the role of FAM162A on the mitochondrial function and cell viability, we performed loss of function experiments in COS7 cells. Those cells were transfected with three different pLVCTH- shFAM162A constructs (siFAM162A #2, #4 and #5) or with the pLVCTH empty vector (Empty) as control. After 48 h post-transfection, the knock-down of FAM162A was confirmed by immunoblot (Fig. 2A). Firstly, cell viability was assessed by the MTT assay and cell mortality by LDH assay. As seen in figure 2B, the siFAM162A cells had a significant 30% reduction in cell viability (*p*<0.05) and 30% increase in mortality (*p*<0.05). Then, mitochondrial function was assessed in terms of membrane potential, oxygen consumption rate and OXPHOS protein expression. Mitochondrial bioenergetics in siFAM162A cells displayed 50% reduction (*p*<0.05) in the mitochondrial membrane potential as evaluated by TMRE staining in non-quenching mode and live cell microscopy (Fig. 2C); and 20% reduction in the basal respiration *(p<*0.05), 30% reduction in the maximal respiration (*p*<0.0001) and 45% reduction in the spare capacity respiration (*p*<0.0001) as determined by the seahorse analysis (Fig. 2D). We wanted to know if decreased respiratory capacity was due to a reduction in the expression levels of OXPHOS proteins. However, no significant differences were found between FAM162A knock-down cells as compared to controls (Fig. 2E).

**Figure 2:**
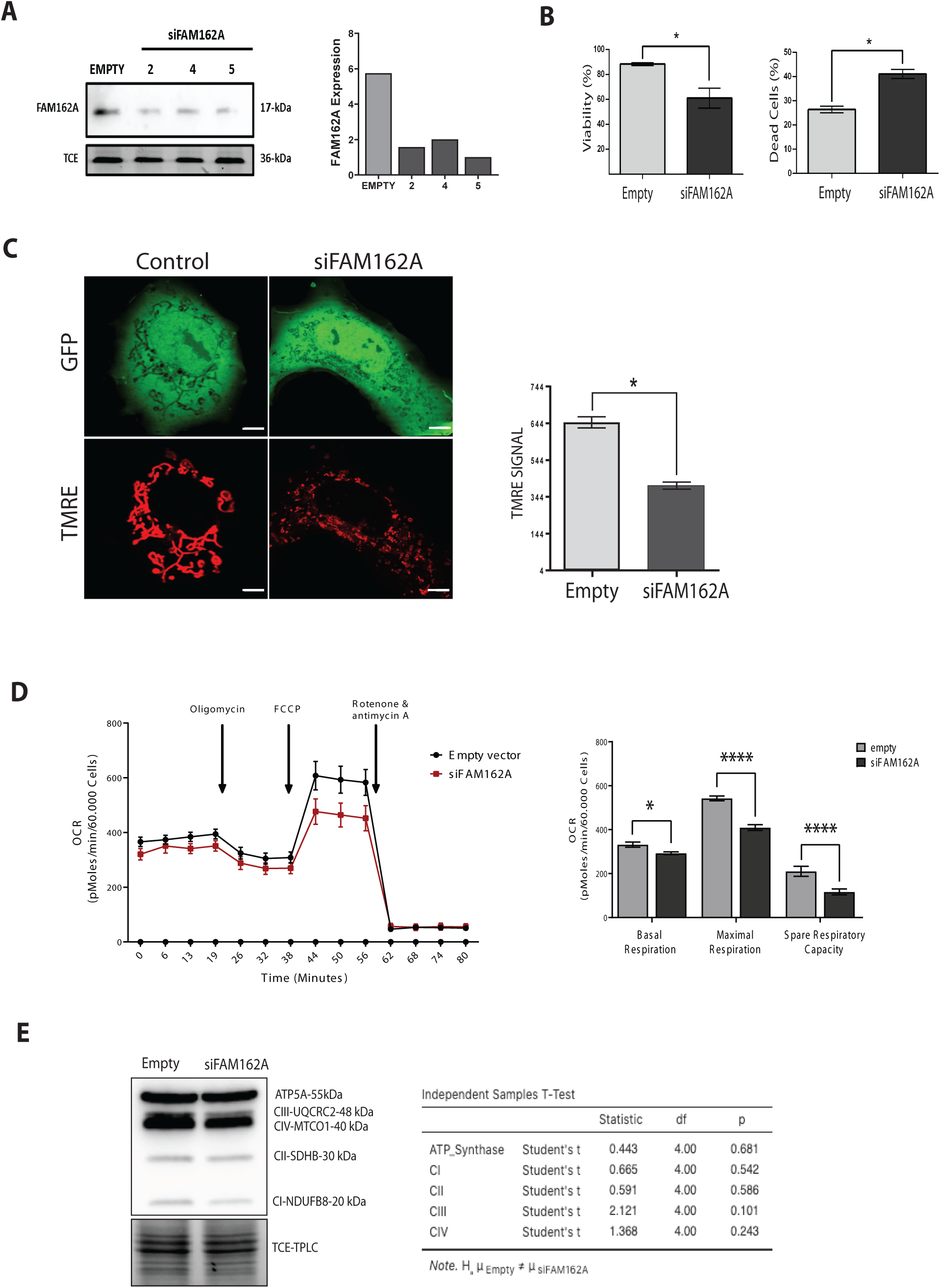
Silencing of FAM162A Disrupts Mitochondrial Bioenergetics and Cell Viability (A) FAM162A Knockdown. Representative Western blot and densitometry quantification showing FAM162A knockdown (siFAM162A) in COS7 cells. Three shRNAs against FAM162A reduced expression by >60%. As loading control, Trichloroethanol (TCE) was included in the gel during casting to visualize proteins *in-membrane* (after the protein transfer) under UV light. **(B) Cell Viability and Cytotoxicity Assays.** MTT and LDH release assays were performed on siFAM162A and control COS7 cells to assess viability and cytotoxicity respectively. Decreased viability and increased mortality was observed in siFAM162A cells as compared to controls. **Statistical analysis, *t*-test, *p*<0.05 (*n*=3).** **(C) Mitochondrial Membrane Potential.** Confocal microscopy on FAM162A knockdown COS7 cells expressing GFP (green channel). Mitochondria were stained with 10nM TMRE (red channel), and the mean fluorescent intensity of TMRE per cell were quantified with the Image J software. Decreased membrane potential was observed in siFAM162A cells as compared to controls. Statistical analysis, *t*-test, *p*<0.05 (*n*=30 cells). **(D) Oxygen Consumption Rate (OCR).** Representative Seahorse assay comparing siFAM162A and control cells. Red line: siFAM162A; Black line: control (Left Panel). Basal, maximal, and spare respiratory rates were measured and quantified (Right Panel) following the sequential addition of oligomycin, FCCP, and rotenone/antimycin A. siFAM162A cells exhibited reduced respiratory rates. Statistical analysis, *t*-test, *p<0.05, ****p<0.0001 (*n*=3). **(E) OXPHOS Protein Expression.** Representative Western blot and quantification of respiratory complexes (CI, CII, CIII and CIV) and ATP synthase in siFAM162A versus control cells. As loading control, Trichloroethanol (TCE) was included in the gel during casting to visualize proteins *in-membrane* (after the protein transfer) under UV light. No significant differences were found. Statistical analysis, *t*-test, *p*>0.05 (*n*=3).

### FAM162A supports mitochondrial morphology and fusion dynamics through OPA1

Qualitative analysis of mitochondrial morphology was performed from TMRE-stained confocal images of siFAM162A and control cells. Mitochondrial units were classified into four categories as described by Leonard et al. (2015) (Leonard et al., 2015): Puncta, large and round, rod (with no branches) and network mitochondria (with branches). FAM162A knockdown caused an increase in puncta mitochondria from 21% to 34%, and a decrease in network mitochondria from 26% to 9% as compared with control (empty) cells (Fig. 3A).

**Figure 3:**
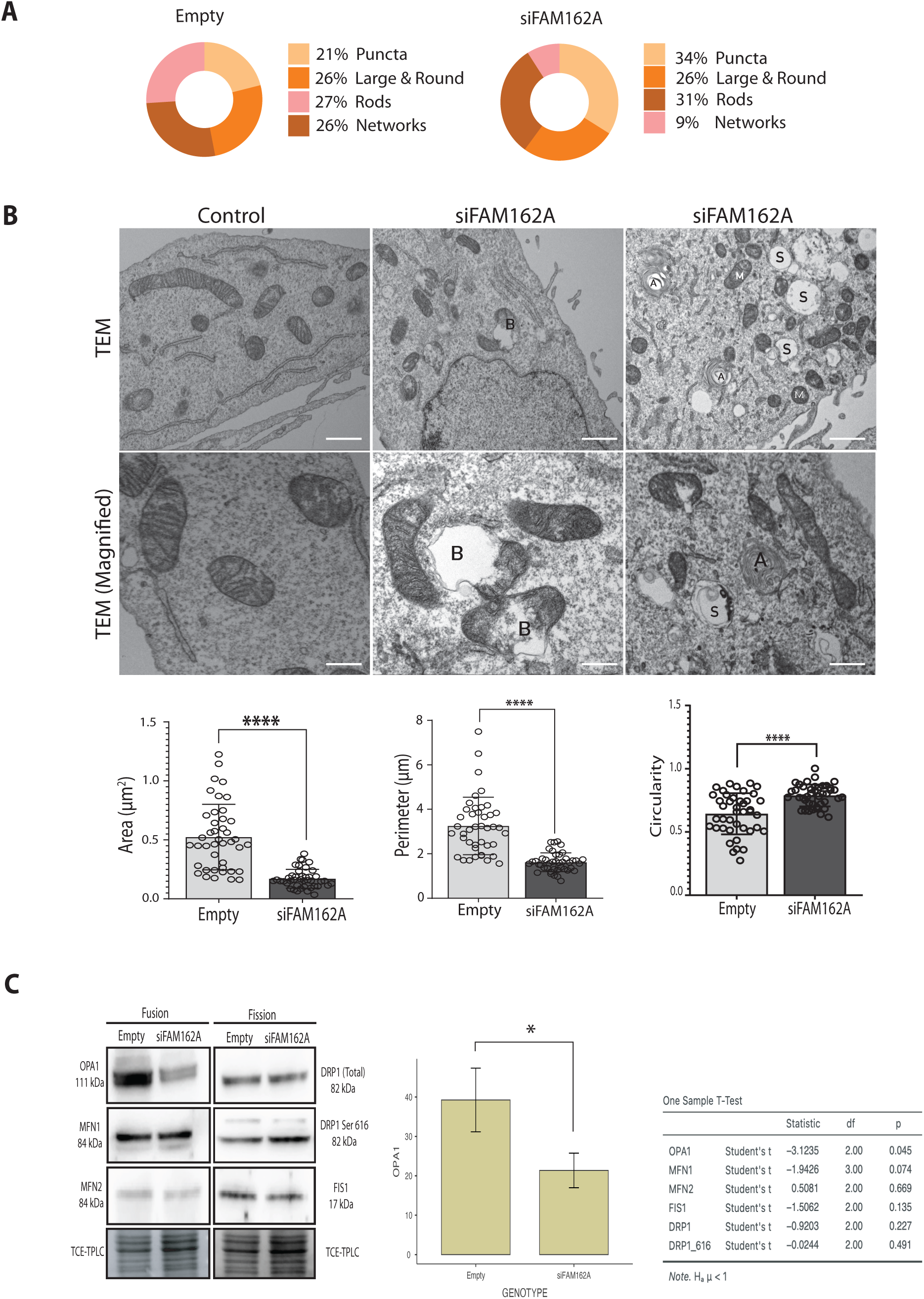
**Loss of FAM612A Disrupts Mitochondrial Morphology and Dynamics (A) Mitochondrial Morphology**. Mitochondrial morphologies were analyzed from TMRE-stained confocal images of siFAM162A and control cells (Fig. 2C) and classified into four categories: Network, Rods, Puncta, and Large & Round. A shift from a network to a punctate morphology was observed in siFAM162A cells (*n*=30) **(B) Mitochondrial Ultrastructure.** Representative TEM images showing mitochondria in control and siFAM162A COS7 cells. Notable features in siFAM162A cells include fragmented mitochondrial morphology (M), bubble-like broken mitochondria (B), swollen mitochondria (S) and an increase number of autophagosome-like structures (A). In addition, mitochondrial area and perimeter were reduced and circularity increased in siFAM162A cells. Scale bar: 1μm or 500nm. Statistical analysis, *t*-test \*\*\*\**p*<0.0001. *n*=80. **(C) Mitochondrial Dynamics Protein Expression.** Representative Western blot and quantitative analysis of mitochondrial fusion proteins (OPA1, MFN1 and MFN2) and mitochondrial fission proteins (DRP1 Phosphorylated DRP1 at Ser616 and FIS1) in siFAM162A cells compared to control. A bar graph showing OPA1 expression highlights a significant reduction in OPA1 levels in siFAM162A cells. As loading control, Trichloroethanol (TCE) was included in the gel during casting to visualize proteins *in-membrane* (after the protein transfer) under UV light. Statistical analysis, *t*-test, p<0.05 (*n*=3).

At the level of ultrastructure, mitochondria from siFAM162A cells were significantly smaller (*p*<0.05) than control mitochondria (Fig. 3B) in terms of area and perimeter. In addition, mitochondrial circularity, where a value of 1 is a perfect circle and 0, not a circle, indicated that siFAM162A mitochondria were significantly rounded as compared with control mitochondria (Fig. 3B). Interestingly, a higher proportion of bubble-like swollen mitochondria was present in siFAM162A cells as compared to controls (Fig. 3B). These mitochondria had broken the outer mitochondrial membrane and presented an evagination of the inner mitochondrial membrane, that was reaching and being exposed to cytosol, and building an electron-lucent single-membrane vesicle, resembling a bubble (Fig. 3B). These mitochondrial bubbles can reach a considerable size up to 2µm diameter. According to Sesso et al. (2012)(Sesso et al., 2012), those bubble-mitochondria correspond to apoptotic mitochondria which were not degraded and therefore accumulated in siFAM162A cells. Furthermore, several swollen mitochondria, autophagosomes and mitochondria-containing autophagosomes were seen siFAM162A cells given to the cell cytosol a vacuolated phenotype (Fig. 3B). Apoptotic and swollen mitochondria in siFAM162A cells may explain the bioenergetic deficiency which may have an impact on cell viability and mortality.

Alterations in mitochondrial morphology prompted us to examine the expression of key mitochondrial dynamics proteins. Notably, we observed a significant reduction (*p*<0.05) in OPA1 levels, which decreased by 50% upon silencing of FAM162A (Fig. 3C). In contrast, the expression of MFN1, MFN2, as well as the fission proteins DRP1, phosphorylated DRP1 (Ser616), and FIS1, remained unchanged. These findings are particularly compelling given FAM162A’s localization in the IMM, suggesting it plays a crucial role in maintaining cristae ultrastructure and promoting mitochondrial fusion through OPA1.

### FAM162A increases the cellular metabolic fitness and mitochondrial turnover giving protection against oxidative stress

To properly investigate the role of FAM162A on mitochondrial function and complement the knock-down experiments, we conducted experiments by FAM162A overexpression. FAM162A cDNA was cloned into the pcDNA 3.1 c-myc plasmid, and COS7 cells were transfected. After 24 hours post-transfection, a Western blot was performed to verify FAM162A overexpression (Fig. 4A). Next, we performed a seahorse bioenergetic assay (Fig. 4B) to measure both the oxygen consumption rate (OCR) for oxidative metabolism, and the extracellular acidification rate (ECAR) for glycolytic metabolism. Results showed a significant (*p*<0.0001) increased OCR at the level of basal, maximal and spare respiration; and decreased ECAR at the level of basal and maximal glycolytic capacity in FAM162A-OE cells as compared to control cells (Fig. 4B and C). These results suggested a metabolic shift towards OXPHOS for cells overexpressing FAM162A. Taking advantage of the Seahorse technology, we also measured the ATP production rate to understand whether ATP is made from glycolytic or oxidative metabolism. As expected, FAM162A overexpression induced a 72% increase (*p*<0.0001) in mitochondrial ATP synthesis and a 33% reduction (*p*<0.01) in glycolytic ATP synthesis (Fig. 4D). This reflects a metabolic shift from glycolysis to OXPHOS.

**Figure 4:**
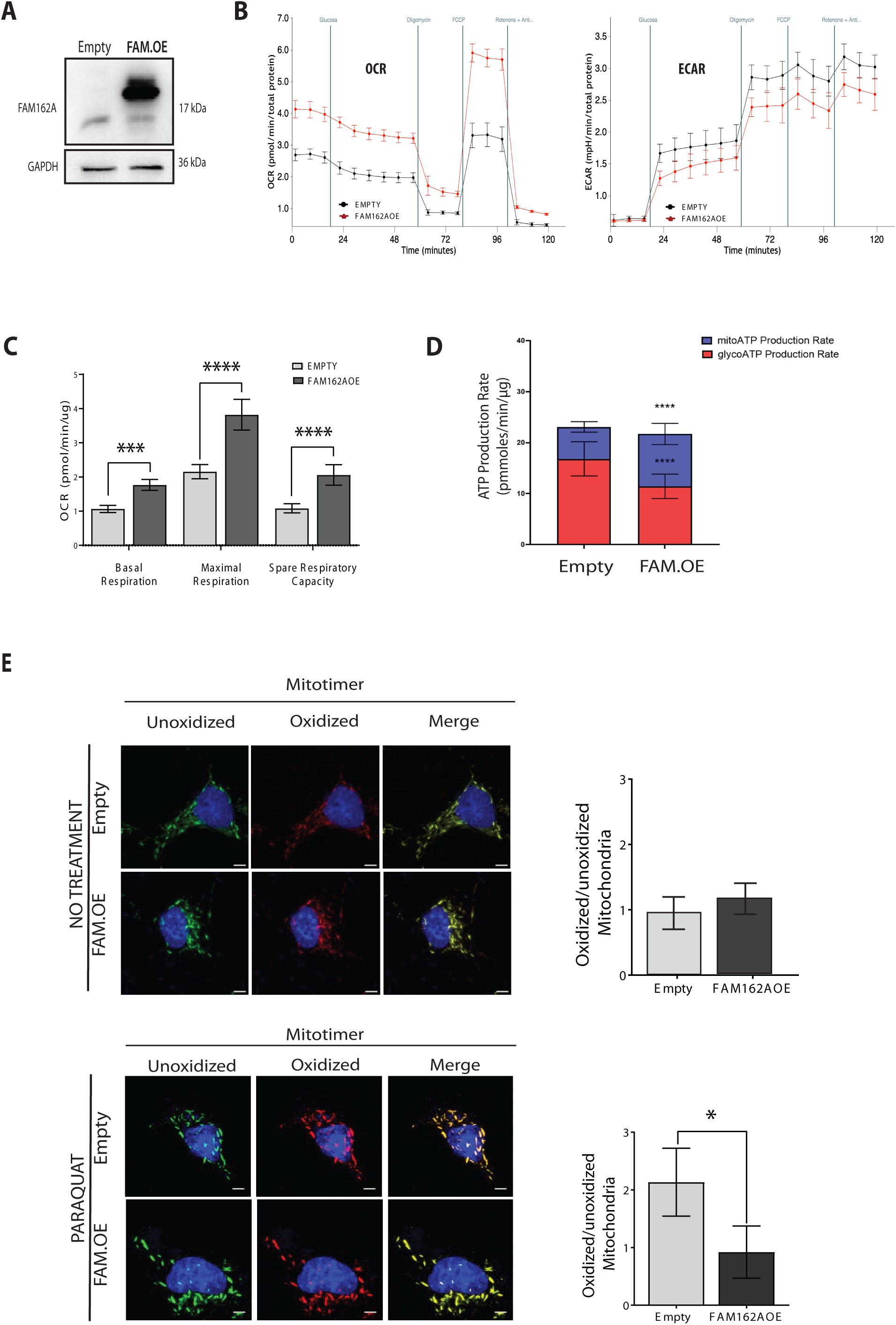
FAM162A overexpression improves mitochondrial bioenergetics and metabolic fitness. (A) Western blot showing FAM162A overexpression (FAM162A_OE) in COS7 cells.(B) Oxygen Consumption Rate (OCR) and Extracellular Acidification Rate (ECAR). A representative seahorse assay is shown for FAM162_OE and control cells. Basal, oligomycin-insensitive, maximal and non-mitochondrial respiration rates were measured as well as basal and maximal glycolytic rate after the sequential addition of oligomycin, FCCP and rotenone/antimycin A. FAM162A_OE displayed higher respiration and lower glycolysis as compared with control cells. Red line: FAM162A_OE cells; Black line: control cells. **(C) OCR quantification.** Seahorse analysis demonstrate a significant increase in basal, maximal, and spare respiratory capacities in FAM162A_OE cells compared to control cells. Statistical analysis, *t*-test, **\***p<0.05, **\*\*\*\***p<0.0001 (*n*=3). **(D) ATP Production Rate in FAM162A_OE and Control Cells.** Using Seahorse technology, both mitochondrial (blue boxes) and glycolytic (red boxes) ATP production rates were measured. Overexpression of FAM162A induced a metabolic shift towards mitochondrial oxidative metabolism. Statistical analysis: t-test, ****p < 0.0001 (*n* = 3). **(E) Mitochondrial Oxidized Protein Levels.** COS7 cells were co-transfected with Mitotimer and FAM162A_OE plasmids, with cells either untreated (upper panel) or treated with 100 µM Paraquat, a pro-oxidant, for 6 hours (lower panel) to induce cellular and mitochondrial oxidative stress. Confocal images were taken at 36 hours post-transfection, capturing Mitotimer fluorescence in both green and red channels. The red (oxidized/old mitochondria) to green (unoxidized/new mitochondria) ratio provides a measure of mitochondrial oxidation. A higher proportion of green mitochondria indicates increased mitochondrial turnover. Under Paraquat treatment, FAM162A-overexpressing cells displayed reduced mitochondrial oxidation, suggesting enhanced mitochondrial turnover. Statistical analysis: t-test, *p < 0.05 (*n* = 3).

To corroborate our results, the mitochondrial health was assessed by co-expressing FAM162A along with the Mitotimer protein (Fig. 4E). Mitotimer is a green fluorescent protein when newly synthesized but transitions to a red fluorescent protein when oxidized over a 42-hour timeframe (Gottlieb and Stotland, 2015; Thornton et al., 2012). COS7 cells were co-transfected to overexpress FAM162A and Mitotimer constitutively and were treated or not with Paraquat (PQ) to induce oxidative damage. Mitotimer’s green (non-oxidized) and red (oxidized) fluorescence were measured at 30 hours post-transfection, and the red/green ratio was calculated as a measure of mitochondrial oxidation. Under no treatment condition, the red/green ratio for FAM162A-OE and control cells was similar with a value close to 1 (Fig.4E, *upper panel*). However, under paraquat treatment, control cell ratio increased to 2.1 ± 0.58 while FAM162A-OE cell ratio remained in 0.9 ± 0.45. This value was significantly lower (*p*<0.05) than the observed in control cells (Fig. 4E, *lower panel*). These Mitotimer results also suggested an increase in mitochondrial turnover when FAM162A is upregulated, protecting mitochondrial function against oxidative stress.

### Transgenic Drosophila Overexpressing FAM162A Have Extended Lifespan and Increased Stress Resistance

Our results suggest that FAM162A contributes to cristae ultrastructure, is associated with mitochondrial fusion, and enhances mitochondrial bioenergetic capacity and turnover. To further investigate FAM162A’s role in vivo, we generated a transgenic fly overexpressing human FAM162A (hFAM162A_OE). The expression of human FAM162A in Drosophila was corroborated by immunoblot (Fig. 5A). As a positive control, COS7 cells overexpressing FAM162A were used, and the UAS_Fam162A control fly served as the negative control.

**Figure 5:**
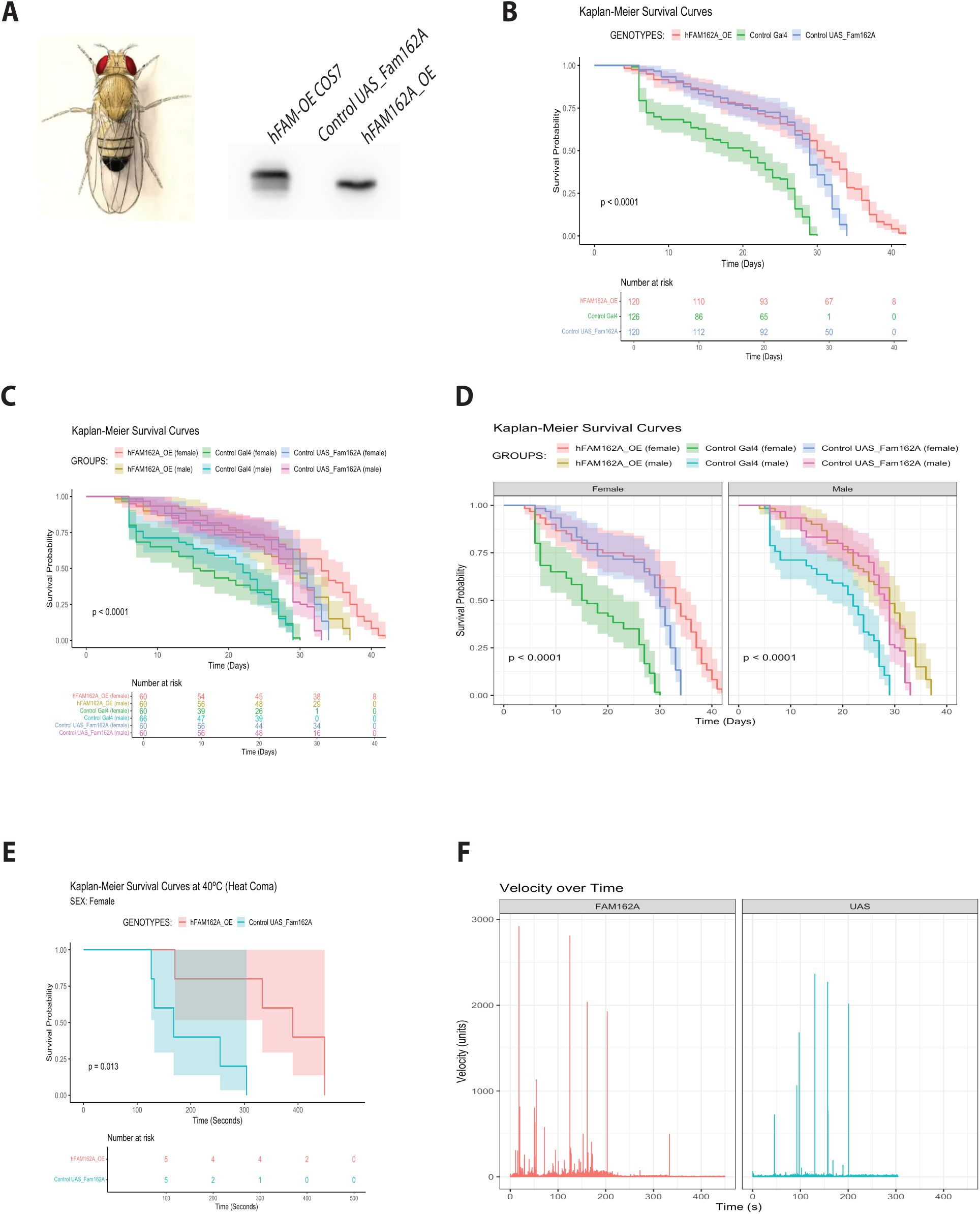
Human FAM162A Overexpression in Transgenic Drosophila (A) FAM162A overexpression in Drosophila. Western blot showing human FAM162A overexpression (hFAM162A_OE) in transgenic Drosophila (genotype Tub-Gal4). FAM162A_OE COS7 cells were used as a positive control. UAS_Fam162A control fly served as the negative control. **(B) Survival Probability of Drosophila.** Survival analysis was conducted on both males and females across three genotypes at 29°C: Control Gal4, Control UAS_Fam162A, and hFAM162A_OE. Kaplan-Meier survival plots were generated using pooled data from both sexes, with a “Number at Risk” table showing live flies for each genotype over time. Overexpression of hFAM162A significantly extended Drosophila lifespan. Statistical analysis: Log-Rank Test; experimental unit: culture tube with 20 flies; *n*=3, ****p<0.0001. **(C) Survival Probability of Drosophila by Sex.** Kaplan-Meier survival plots and “Number at Risk” tables are shown for each genotype at 29°C, separated by sex. Female Drosophila overexpressing hFAM162A demonstrated a significantly longer lifespan compared to males. Statistical analysis: Log-Rank Test; experimental unit: culture tube with 20 flies; *n*=3, ****p<0.0001. **(D) Survival Probability by Sex, faceted view.** This panel provides Kaplan-Meier survival curves for each sex individually to better highlight the lifespan differences in hFAM162A_OE flies compared to controls. **(E) Survival Probability of Female Drosophila under Heat Stress.** Survival analysis was conducted on female Control UAS_Fam162A and hFAM162A_OE flies at 40°C, with time recorded in seconds. Kaplan-Meier survival plots and “Number at Risk” tables demonstrate an extended lifespan for hFAM162A-overexpressing females under heat stress. Statistical analysis: Log-Rank Test; experimental unit: culture tube with 1 fly; *n*=5, ****p<0.0001. **(F) Locomotor Activity of Female Drosophila under Heat Stress.** Activity, measured as velocity adjusted by weight, was recorded until fly death and analyzed using Bonsai tracking software. Only the first 100 seconds were analyzed (when all flies were still alive, per the “Number at Risk” table in panel E). hFAM162A_OE flies showed greater locomotor activity at 40°C compared to controls. Experimental unit: culture tube with 1 fly; *n*=5.

For lifespan assessment, twenty flies of each genotype, separated by sex (male and female), were housed in fly-culture tubes at 29°C. Survival was tracked over time, recording the time of death for each fly. This experiment was repeated three times using flies from independent crosses. At 29°C, pooled lifespan data indicated that flies overexpressing human FAM162A showed a significant increase in survival (*p*< 0.001) compared to both Control-Gal4 and Control UAS_FAM162A flies, as determined by the log-rank test with pairwise comparisons (Fig. 5B). Differences in survival between hFAM162A_OE and Control UAS-FAM162A became apparent around day 30, with hFAM162A_OE flies exhibiting a 25% longer lifespan overall. When survival data was separated by sex, we found that FAM162A overexpression had a more pronounced effect in females, extending their lifespan by approximately 12.5% more than males (*p*<0.001, Fig. 5C). For clarity, sex-specific survival plots are shown in Fig. 5D, indicating an inflection point around day 30 for both sexes at 29°C.

To evaluate stress resistance, transgenic flies were exposed to heat stress at 40°C, during which lifespan and locomotor activity (measured as velocity adjusted by weight) were assessed. Under heat stress, hFAM162A_OE flies survived a significantly longer 40% and displayed enhanced heat stress resistance (*p*<0.05) than controls. At 300 seconds, four out of five hFAM162A_OE flies remained alive, compared to only one out of five for Control UAS_FAM162A (Fig. 5E). Furthermore, hFAM162A_OE flies exhibited increased locomotor activity as compared to control flies (Fig. 5F). This trend held true even during the first 100 seconds, when all flies were still alive, reflecting the superior metabolic capacity of hFAM162A_OE transgenic flies.

## Discussion

Mitochondria plays a pivotal role in cell differentiation, proliferation, and survival. Conversely, dysfunctional mitochondria have been implicated in aging, cancer and neurodegenerative and cardiovascular diseases among others. Intriguingly, a healthy mitochondria population has the potential to restore cell function and reverse associated symptoms, as evidenced in both cellular and animal models. This underscores the medical significance of exploring novel mitochondrial proteins and their regulatory mechanisms (Elorza and Soffia, 2021; Schmid et al., 2022; Gumeni et al., 2021; Doblado et al., 2021).

In this research, we get a deeper understanding of the hypoxia-induced apoptotic protein FAM162A (Lee et al., 2004), a membrane bound mitochondrial protein. We found that FAM162A plays a role in preserving the mitochondrial ultrastructure and bioenergetics. Loss of function experiments performed in COS7 cells showed that mitochondrial morphology was fragmented with reduced bioenergetic capacity impacting cell viability with increased cell death. At the level of ultrastructure, besides to observe evident mitochondrial fragmentation with reduced size, many other mitochondria were found either enclosed in autophagosomes or in a pre-apoptotic state displaying a bubble-like and swollen morphology. Mitochondrial fragmentation may be due to increased fission or decreased fusion dynamics. Our results suggested a reduction in mitochondrial fusion especially due to decreased protein levels of OPA1. When the membrane topology and submitochondrial localization of FAM162A was interrogated, which had not been previously studied, we determined that FAM162A is in the inner mitochondrial membrane, mainly in the cristae membrane, with both ends facing the mitochondrial matrix. This finding aligned with the reduced OPA1 levels as OPA1 is also localized in the cristae membrane, suggesting a potential interaction, direct or indirect, between FAM162A and OPA1.

On the other hand, FAM612A overexpression increased both the bioenergetic capacity promoting a metabolic switch from glycolysis to OXPHOS metabolism, and mitochondrial turnover, making cells more resistant to oxidative stress. Most of these results were in accordance with our hypothesis, unveiling the importance of FAM162A for mitochondrial function. Considering the pronounced phenotypic effects observed upon overexpression or silencing of FAM162A on mitochondrial structure, dynamics, bioenergetics, and its localization within cristae, questions about the underlying cellular and molecular mechanisms arise.

Previous studies have reported that FAM162A directly interacts with VDAC, a protein located in the OMM (Lee et al., 2004). VDAC interacts with the “mitochondrial contact site and cristae-organizing system” (MICOS) complex, implicated in maintaining cristae structure and cristae junctions crucial for oxidative phosphorylation (OXPHOS) and ATP synthesis (Anand et al., 2021; Yang et al., 2022; Colina Tenorio et al., 2020). The cristae junction acts as a bridge that allows communication and exchange of molecules between the cristae and the rest of the mitochondria (Joubert and Puff, 2021). Importantly, key proteins involved in cristae organization besides MICOS include the fusion protein OPA1, and dimers of the ATP synthase (Cogliati et al., 2013; Stroud and Ryan, 2013). OPA1, besides to acts as a master regulator of mitochondrial fusion (Song et al., 2009), plays a crucial role in maintaining the integrity and remodeling of cristae and is essential for the assembly of respiratory complexes by means of OPA1-OPA1 and OPA1-MICOS interactions (Kondadi et al., 2020; Anand et al., 2021; Hu et al., 2020; Jang and Javadov, 2020). The ATP synthases form rows of dimers which induce membrane curvature to give cristae lamellar or tubular morphology to maximize the use of ΔΨm for ATP production. The interplay between MICOS, OPA1 and ATP synthase is involved in shaping cristae. Cristae undergo dynamic changes in number and shape, adapting to energy demands and can also operate as energetically independent units (Miranda-Astudillo et al., 2022; Cadena et al., 2021; Wolf et al., 2019). Dysregulation or mutations in MICOS, OPA1, and ATP synthase can impair cristae structure, function, and the assembly of OXPHOS, resulting in various human diseases associated with neurodegeneration and mitochondrial dysfunction. In this regard, FAM162A may also be part of the regulatory system of the cristae junction and structure.

Additionally, VDAC participates in the regulation of mitophagy, acting as a pivotal factor in determining whether a mitochondrion undergoes apoptosis or mitophagy based on its pattern of ubiquitination, which is generated by PARKIN (Ham et al., 2020; Ordureau et al., 2018). Several lines of evidence suggest that VDAC is involved in PINK1/Parkin-mediated mitophagy, facilitating the recruitment of PARKIN from the cytosol to mitochondria in various cell types (Geisler et al., 2010; Sun et al., 2012; Yang et al., 2021; Xie et al., 2021). Interestingly, cellular models of Parkinson’s disease generated due to OXPHOS inhibition at complex I, displayed both mitochondrial fragmentation and enlargement, dysregulation of autophagy due to decreased PINK1and PARKIN protein levels; and a significant reduction in FAM162A protein levels (Mazzio and Soliman, 2012; Zhu et al., 2013; Verma et al., 2020; Ma et al., 2020). Based on these observations, and given that FAM162A induced mitochondrial turnover, especially under stress conditions, we hypothesized the existence of a FAM162A-OPA1-VDAC axis that regulates mitochondrial architecture and bioenergetics, possible linked to mitophagy. Interestingly, elevated FAM162A protein levels have been observed in various cancer types, correlating with enhanced proliferation and migration capacities. In fact, not only were FAM162A levels increased, but also the mitophagic proteins NIX and BNIP3 (Tang et al., 2009; Cho et al., 2010; Nissou et al., 2013; Sorensen et al., 2015; Toustrup et al., 2011) The generalizability of our cellular and molecular results was limited by the fact that those experiments were done on COS7 cell line which is derived from monkey’s kidneys. Therefore, we wanted to explore the role of FAM162A in an organism model to assess its universal function *in vivo*. As outlined in the results section, FAM162A exhibits broad expression across various cell types and taxa, suggesting its significance in cellular and organismal physiology. To this end, we generated the transgenic Drosophila overexpressing human FAM162A and observed, both in males and specially in females, that lifespan and locomotor activity were increased under normal and heat stress conditions. Stress resistance has been linked to longevity in various organisms. In Drosophila, it has been shown that longer-lived females had lower ROS levels and higher superoxide dismutase and catalase activity, both antioxidant enzymes, as compared with males. Furthermore, under ethanol stress, females showed greater resistance to mortality and better locomotor function (Niveditha et al., 2017). Similar results have been found in studies involving mitochondrial and mitophagy-related proteins. When mitochondrial dynamics is stimulated by DRP1 overexpression, a mitochondrial fission protein, females had an extended lifespan and improved physical activity in midlife, alongside enhanced stress resistance, compared to males (Rana et al., 2017). On the other hand, mutations in Drosophila OPA1-like, besides having neurological defects, presented reduced lifespan (McQuibban et al., 2006). When assessing autophagy pathways, it has been reported that the overexpression of PINK1, PARKIN and P62 prolonged lifespan in Drosophila due to increased mitochondrial proteostasis and improved mitochondrial function (Si et al., 2019; Aparicio et al., 2019).

In summary, our study identifies FAM162A as a novel gatekeeper of mitochondrial function contributing to mitochondrial integrity, bioenergetics, mitochondrial dynamics and turnover, providing stress resistance and longevity both in cellular and Drosophila models.

**Declarations section**

## Ethics Approval and Consent to participate

Not applicable.

## Consent for publication

Not applicable.

## Availability of data and materials

The datasets used and/or analyzed during the current study are available from the corresponding author on reasonable request.

## Competing interests

The author(s) declare(s) that there is no conflict of interest regarding the publication of this article.

## Funding

This research was supported by the National Agency for Research and Development (ANID) [FONDECYT grant number 1180983 to A. A. Elorza; FONDEQUIP grant number EQM220115 to A. A. Elorza; Ph.D. Scholarship, number 2021-21212271 to A. Matamoros]; the National Institute of Health [Research Project grant number 5RO1S108778 to R. Jorquera]; and the Universidad Andres Bello [Nucleus-UNAB grant number DI-03-22/NUC to A. A. Elorza; Research Initiation Fellowship DGI-UNAB grant number DI-13-22/INI to A. Matamoros, and grant number DI-07-22/INI to G. Gomez-Lillo; Ph.D. Scholarships to both J.P. Soffia and G. Gomez-Lillo].

## Author Contributions

**A. Matamoros** conducted most of the experiments, analyzed data, make figures, partially funded the research, and help to write and review the manuscript. **J.P. Soffia** conducted Drosophila experiments and analyzed fly data. **M. Maturana** performed the localization, topology and orientation experiments of FAM162A. **M. Muñoz** conducted the bioenergetics experiments for the siFAM162A cells. **A. Gonzalez-Ibañez** performed bioinformatics analysis and supported biochemical and microscopy experiments. **G. Gomez-Lillo** supported biochemical and microscopy experiments. **C. Astorga** designed Drosophila vectors and transgenics fly and supported Drosophila experiments. **L. M. Ruiz** performed the TEM experiments. **R. Jorquera** supported Drosophila experimental design, analyzed fly data and funded fly transgenic generation. **A. A. Elorza** conceived the idea, designed the experiments, analyzed data, wrote the manuscript, make figures, and funded the research.

## Declaration of Generative AI and AI-assisted technologies in the writing process

Statement: During the preparation of this work the author(s) used ChatGPT partially to improve English language for clarity and conciseness. After using this tool/service, the author(s) reviewed and edited the content as needed and take(s) full responsibility for the content of the publication.

## Acknowledgments

We thank Dr. Roberta Gottlieb (Cedars-Sinai Medical Center, CA, USA) and Dr. Orian Shirihai (UCLA, CA, USA) for their thoughtful insights, comments, and discussions about this work.

## Notes

### Competing Interest Statement

The authors have declared no competing interest.

